# Seeing emotions, reading emotions: behavioral and ERPs evidence of the effect of strategy and of regulation for pictures and words

**DOI:** 10.1101/490409

**Authors:** Alessandro Grecucci, Simone Sulpizio, Elisa Tommasello, Francesco Vespignani, Remo Job

## Abstract

**Background:** Whilst there has been extensive study of the mechanisms underlying the effect of regulation for the emotions elicited by pictures, the ability and the mechanisms beyond the regulation of words remains to be clarified. Similarly, the effect of strategy when applying a regulatory process is still poorly explored. The present study seeks to elucidate these issues comparing the effect of regulation and of strategy to both neutral and emotional words and pictures.

**Methodology/Principal Findings:** Thirty young adults observed and took the distance from unpleasant and neutral pictures and words while their subjective ratings and ERPs were recorded. At a behavioral level, participants successfully regulated the arousal and the valence of both pictures and words. At a neural level, unpleasant pictures produced an increase in the late positive potential modulated during the regulate condition. Unpleasant linguistic stimuli elicited a posterior negativity as compared to neutral stimuli, but no effect of regulation on ERP was detectable. More importantly, the effect of strategy independently of stimulus type, produced a significant larger Stimulus Preceding Negativity. Dipole reconstruction localized this effect in the middle frontal areas of the brain.

**Conclusions:** As such, these new psychophysiological findings might help to understand how pictures and words can be regulated by distancing in daily life and clinical contexts, and the neural bases of the effect of strategy for which we suggest an integrative model.

## Introduction

Visual, auditory and other sensory modalities have specific neural circuitry to process the emotional meaning carried by the incoming stimuli. Hardwired in specific brain mechanisms such a circuitry evolved to respond to relevant situations with efficient responses. However, for successful adaptation to an ever-changing environment, sometimes in-coming stimuli need to be regulated before being channeled into action [1,2]. This may be necessary in order not to be overwhelmed or distracted by the emotional impact of the situations. Although in the last twenty years we have come to know the basic neural structures involved in emotional perception and regulation by means of the extensive usage of neuroimaging techniques such as functional Magnetic Resonance Imaging (fMRI) (see [3], for a review), the temporal dynamics associated with the regulation of emotional stimuli are poorly understood. In the present paper, we start to overcome this limitation by unveiling electrophysiological activity of people regulating their response to emotional pictures and words. Electroencephalography is possibly the best way to capture the temporal dynamics of complex psychological events that unfold over time since it allows to identify, with a precision of milliseconds, close but subsequent events and to track their temporal development.

A challenge in the actual emotion regulation field would be to investigate the regulation of emotion elicited by words, and to compare such findings with those elicited by pictures. In daily life, we usually express emotions by means of words, and we react with emotions to other people words. A case in point, is what happens during a psychotherapy session where the clinician uses language to elicit and regulate patients’ emotions, and to understand the emotional states of patients by listening to their words [4,1,2]. Freud defined psychotherapy “a cure through words” [5]: a cure to help patients express and possibly regulate linguistic contents. Despite its relevance in daily life and in clinical situations, the experimental investigation of the processes involved in regulating emotions conveyed by words are scant. Indeed, most of the previous experiments were focused on the regulation of emotional pictures rather than emotional words. To our knowledge there are only a few experiments testing the effect of emotion regulation on words. In one experiment [6], participants were asked to regulate emotions to unpleasant pictures and then were asked to judge whether a word was negative or neutral. Event-related potentials (ERPs) revealed a reduction in N400 and an increase P300 amplitudes to words presented after enhancing and suppressing unpleasant emotions, respectively. However, in this experiment the modulation of words followed a previous modulation of pictures, making it hard to disentangle the regulation of pictures from the regulation of words. Moreover, participants performed a decision task while reading the words, and this may have distracted participants from the regulation task. More recently, Speed et al., [7] used an Autobiographical emotion regulation task (AERT) to study the possibility of regulating the emotions elicited by linguistically cued autobiographical memories. However, in this experiment words where used only to cue autobiographical memories, and not as a target of regulation. Thus, the first aim of this paper is to provide clear evidence on the possibility to regulate emotional linguistic stimuli. In order to allow for possible comparisons with well-established findings of effects of regulation, we developed an integrated paradigm in which in separate blocks particpants regulate pictures or words laden emotional stimuli. In each block neutral stimuli were also present.

The majority of previous studies used reappraisal (in the form of reinterpretation) and distraction, two largely studied regulatory strategies [8,9,10]. However, reinterpretation and distraction are just two examples of a vast set of regulation strategies people use. Reappraisal typically involves an effort to generate an alternative interpretation of the incoming stimulus [see, e.g., 11]. Distraction-based strategies, as the one used by Schönfelder and colleagues [12] required subjects to apply mental operations (performing arithmetic calculations while observing the emotional stimuli). In the present study we used distancing [13,14]. This strategy does not require participants to perform inferential (reappraisal) or mathematical (distraction) operations. It simply implies to put oneself in a detached perspective as if the incoming stimulus has nothing or little to do with oneself. Thus, departing from previous studies, the second aim of the study is to test the possibility to regulate emotional words and pictures by using distancing.

Beside the type of strategy used, another challenge in the current debate concerning emotion regulation is the possibility to disentangle the *effect of strategy* from the *effect of regulation*. The effect of strategy (ES) concerns the implementation of the strategy while the effect of regulation (ER) is related to the regulation of emotions to which the strategy was applied. In a functional magnetic study, Grecucci et al [15,16] showed that different brain networks were linked to ES (how the brain responds when the subjects applied the strategy) and to ER (how specific brain regions are regulated by the strategy). Specifically, strong activation of dorsolateral prefrontal cortex was associated with ES while reduced activation of insula, among other structures, was associated with ER. These results were based on statistical contrasts inside the same temporal window, thus making it hard to disentangle the separate effects of the two processes. More related to EEG, authors found in the reappraisal condition a reduced Late Positive Potential (LPP) as possible ER. LPP is a positive slow component with a posterior distribution, which is sensitive to the emotional content of the stimuli and is larger for emotional than neutral stimuli [e.g., 17]. A reduction of LPP was also reported by Moser et al. [18], with the strategy suppress, and by Foti & Hajcak [19] who investigated the modulation of neutral vs negative effect of descriptions before the image was presented. Schönfelder and colleagues [12] showed a reduction in emotional activation when participants used a distraction strategy but also, to a lesser extent, when they used a reappraisal-based strategy. Paul et al., [20] showed a reduction of LPP when participants used distraction and reappraisal-based strategies. Gan and colleagues [21] replicated the effect of reduction of emotion enhanced LPP when participants applied reappraisal and observed an increase of N2 for the suppression strategy, possibly related to the control of facial expression. More recently, Qi et al., [22], used detached reappraisal (similar to what we indicate with the term distancing) to regulate unpleasant pictures and reported a modulation of LPP. However, these experiments mainly focused on the ER. Studies that specifically focused on the ES found an increased Stimulus Preceding Negativity [SPN, 23-25, but see also 26 for null results] in the pre-implementation period, for reappraisal but not for distraction [24], and interpreted it as enhanced recruitment of attentional resources. In the present experiment, we aimed at temporally separating ES from ER by detecting the associated ERPs in their respective temporal windows (aim 3). In order to do so, we analyzed two well defined and distant time windows, to detect separate cortical responses of the ER from the ES. This can lead to the discovery of specific neural markers when participants apply the strategy of distancing (ES) beside effects of emotion regulation (ER). In sum, we aim at testing three main hypotheses: 1) we hypothesize that emotional words can be regulated in both the arousal and valence dimensions in a similar fashion to what has been previously showed for pictures; 2) we hypothesize that distancing is able to regulate both emotional words and pictures; last but not least 3) we hypothesize that separate and different ERP components can be associated with the effect of regulation (in the time window of the stimulus to regulate) from the effect of strategy (in the time window of the presentation of the strategy to apply before the stimulus presentation). The effect of regulation should be manifested in well-known components such as the modulation of LPP. By contrast, we hypothesize a different component at anterior sites for the effect of regulation, based on previous EEG [23-25], and fMRI studies [15,16].

## Method

### Participants

Thirty-one adults participated in the Experiment. The data of one participant were discarded because of the very high number of artifacts in the EEG data. The final sample was of 30 participants (23 female, mean age: 22.37, SD: 3.23; mean years of education: 15.93, SD: 2.63). All participants were right-handed, Italian native speakers; they had normal or corrected-to-normal vision, and reported to be neurologically healthy. Participants gave written informed consent to their participation after they were informed about the nature of the study. Participants received a small financial reward to take part to the study. The study was approved by the ethical Committee of the University of Trento.

## Materials and design

### Behavioral paradigm

The experiment comprised both pictures and words as stimuli which participants were trained to attend (baseline condition), or to regulate (experimental condition) upon. Pictures and words stimuli were taken from the International Affective Picture System (IAPS, [27]), and from the Affective Norms for English Words (ANEW for Italian, [28]), respectively. These stimuli could be neutral and negative according to their valence. Eighty stimuli per each category were selected for a total of 160 pictures and 160 words. The negative pictures had low valence (M=2.56) and medium-high arousal (M=6.09). The neutral pictures had medium valence (M=5.29) and low arousal (M=3.09). Both neutral and negative pictures were divided into two subsamples and associated with the two experimental conditions (attend vs distancing). Importantly, they did not statistically differed (valence of negative subsample 1 vs negative subsample 2, t (78) = 0.68, p = .946; arousal of negative subsample 1 vs negative subsample 2, t (78) = -0.257, p = .798; valence of neutral subsample 1 vs neutral subsample 2, t (78) = -0.787, p = .434; arousal of neutral subsample 1 vs neutral subsample 2, t(78) = 0.277, p = .783).

The negative words had low valence (*MEAN* = 2.42) and medium-high arousal (*MEAN* = 6.45). The neutral words had medium valence (*MEAN* = 6.07) and low arousal (*MEAN* = 4.15). Negative and neutral words were matched on the following psycholinguistic variables: Written frequency, length in letter, and orthographic neighborhood size (t < 1.68, p >.09). Stimuli were divided into two subsets (one per experimental condition). Every participant was exposed to one of the two subsets associated with one of the two experimental conditions. Both neutral and negative words were divided into two subsamples and associated with the two experimental conditions (attend vs distancing). They did not statistically differed (valence of negative subsample 1 vs negative subsample 2, t (78) = -0.296, p = .768; arousal of negative subsample 1 vs negative subsample 2, t (78) = -0.823, p = .413; valence of neutral subsample 1 vs neutral subsample 2, t (78) = 0.505, p=.615; arousal of neutral subsample 1 vs neutral subsample 2, t (78) = 0.223, p = .824).

After the fixation point (1000 ms), instructions on the regulatory strategies were given (attend, or distancing, for 3000 ms). This was the time window to extract the signal relative to the Effect of Strategy. To avoid, linguistic confounding we used a circle to indicate the attend condition, and a downward arrow to indicate distancing. Then the stimulus (picture or word separated in blocks) was presented for 1000 ms. After the stimulus, a black screen appeared for 4000 ms. In sum, 5000 ms were given for the perception and regulation of emotions elicited by the stimuli. Then participants rated their emotions according to the valence and arousal dimensions on two separate scales, by using the Self-Assessment Manikin (SAM) procedure on a 9-points Likert scale [29]. An interstimulus interval of 3000 ms was then presented before next trial. See Figure 1A.

**Figure 1.**
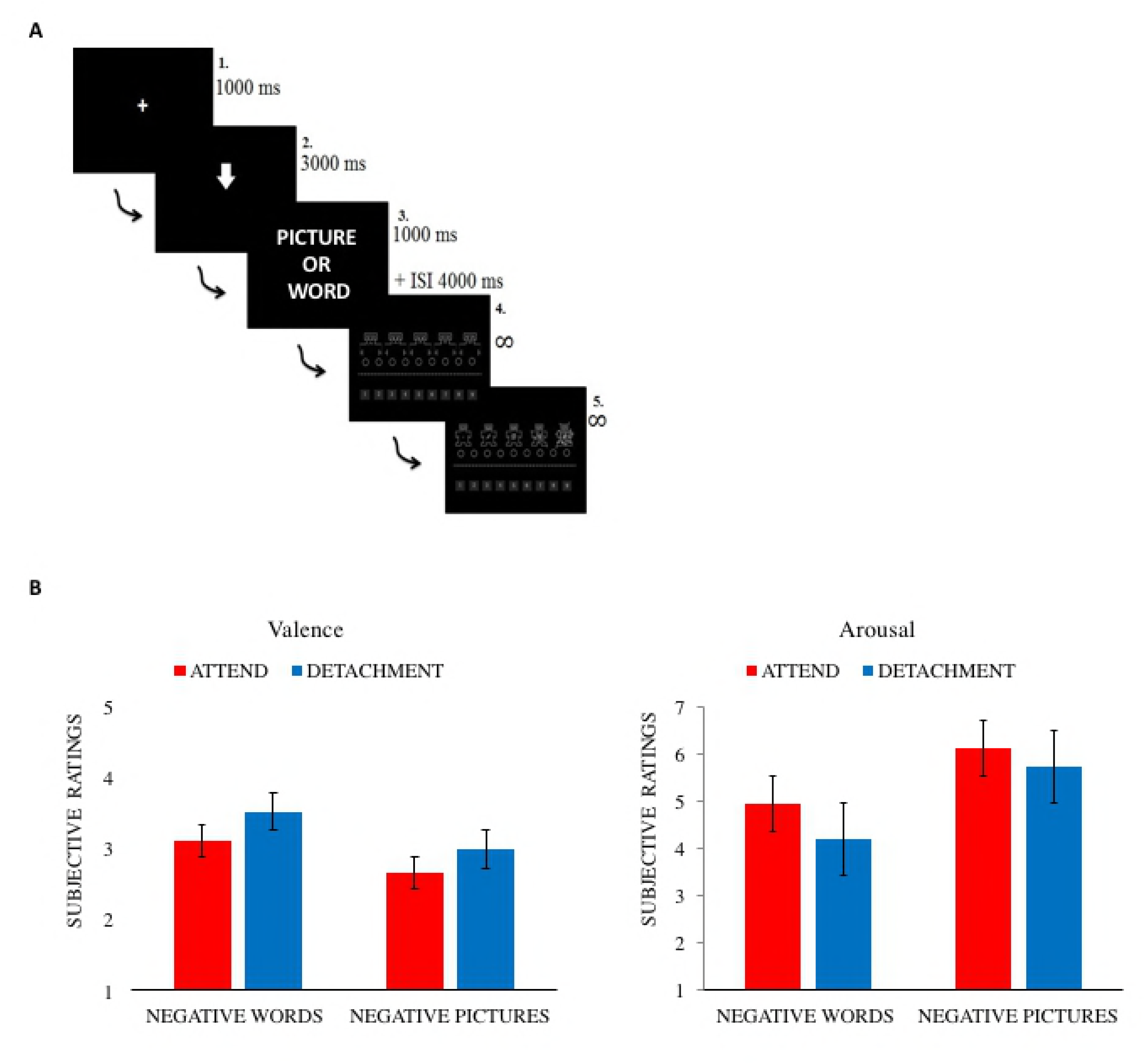
Timeline of events (A) and behavioral results (B). Participants showed an effect of regulation for both words and pictures when applying the regulation strategy. Distancing reduced arousal (strength of perceived emotions), and increased valence (perceived as less negative) for negative pictures and words. Error bars indicate the standard error of the mean, asterisks significant differences.

Before the experiment, a written protocol describing how to apply the emotion regulation strategy of distancing was given. In brief, in this protocol, participants were asked to put themselves in a detached perspective as if that event was far from their lives and not connected at all with them. Two examples were given. A negative picture, and a negative word were presented accompanied by the explanation on how to apply the strategy. In the attend condition, they were asked to respond naturally, without using any specific strategy.

### EEG Procedure

EEG was recorded from 60 scalp electrodes (Fp1, Fpz, Fp2, AF7, AF3 AF4, AF8, F9, F7, F5, F3, F1, Fz, F2, F4, F6, F8, F10, FT7, FC5, FC3, FC1, FCz, FC2, FC4, FC6, FT8, T7, C5, C3, C1, Cz, C2, C4, C6, T8, TP7, CP5, CP3, CP1, CPz, CP2, CP4, CP6, TP8, TP10, P7, P5, P3, P1, Pz, P2, P4, P6, P8, PO7, PO3, POz, PO4, PO4, PO8, O1, Oz, O2) mounted on an elastic cap, positioned according to the international standard position (10-20 system). Additional external electrodes were placed 2 below the eyes (Ve1, Ve2) and 2 lateral to the external canthi (He1, H2). Electrodes were referenced to Cz; the ground was placed anteriorly to AFz. Impedance was kept below 10 kΩ. Data were acquired at the sampling rate of 1000 Hz with a low-pass filter with 100 Hz cutoff frequency and 10s time constant.

Offline analysis was performed with Brain Vision Analyzer II software (Brain Products GmbH, Munich, Germany). Data were down sampled at 250 Hz and re-referenced to an average reference; the signal of Cz was reconstructed. Data were filtered with a low pass filter (40 Hz cutoff, 12 dB/oct) and a high-pass filter (0.05 Hz cutoff, 12 dB/oct sec). Two virtual EOG channels were off-line computed as the difference between the average of Fp1 and Fp2 and the average of Ve1 and Ve2 (VEOG), and as the difference between He1 and He2 (HEOG). The signal was corrected for eye blinks and ocular movements by means of Independent Component Analysis. Ocular electrodes were then excluded from subsequent analyses as well as other four electrodes (AF7, AF8, F7, F8) due to their very high level of noise in all participants.

The EEG signal was segmented in two ways in order to have strategy segmented epochs, lasting from 500 ms before until 3000 ms after strategy onset, and target segmented epochs, lasting from 500 ms before until 2000 ms after target onset. In both cases, the segmented epochs in which the voltage exceeded [-100μV, 100μV] in any of the channels were rejected as affected by artifacts (overall, the 70% of strategy trials and 77% of target trials were kept for analyses). For the strategy epochs, single subject waveforms for each condition were averaged in reference to the 400 ms prestimulus baseline whereas for the target epochs they were averaged in reference to 300 ms prestimulus baseline.

## Results

### Behavioral results

We first computed a general ANOVA on SAM ratings with all factors: Index (Valence vs Arousal), Regulation (Distancing vs Attend), Stimulus (Pictures vs Words) and Emotional content (Negative vs Neutral). Analysis returned a significant main effect of Regulation (*F*(1,29) = 20.040, p < 0.05), Stimulus (*F*(1,29)= 6.81, p < 0.05), and Emotional content (F(1,29)=4.99, p <0.05), as well as a significant interaction between Index and Regulation (F(1,29)=16.30, p <0.05); Index and Emotional content (F(1,29)=303.65, p <0.05); Stimulus and Emotional content (F(1,29)=42.39, p <0.05); Index, Regulation and Emotional content (F(1,29)=20.46, p <0.05); Index, Stimulus and Type (F(1,29)=41.83, p <0.05).

Next, we computed two separate ANOVAs one for each Stimulus type. For Pictures, analysis returned a significant effect of Regulation (F(1,29)=7.68, p <0.05), Emotional content (F(1,29)=18.09, p <0.05), as well as the interaction between Index and Strategy (F(1,29)=7.84, p <0.05), Index and Emotional content (F(1,29)=235.20, p <0.05), and the triple interaction Index, Regulation and Emotional content (F(1,29)=12.35, p <0.05). To explore the triple interaction, we computed Bonferroni corrected post hoc on the arousal and the valence of each category. For Arousal, there was a significant difference between distancing and attend for negative pictures (p<0.005), but not for neutral pictures (p=0.157). Distancing reduced arousal (strength of perceived emotions) when applied to negative pictures. Neutral and negative pictures differed from each other in both conditions (p<0.001). See Table 1 and Figure 1B. For Valence, there was a significant difference between distancing and attend for both negative pictures (p<0.01), and neutral pictures (p<0.001). Distancing increased valence (perceived as less negative). Neutral and negative pictures differed from each other in both conditions (p<0.0001). See Table 1 and Figure 1B.

**Table 1.** SAM mean ratings and standard deviations for pictures and words.

|  |  |  |  |  |
| --- | --- | --- | --- | --- |
|  | **Pictures** |  |  |  |
|  | **Look** |  | **Distancing** |  |
|  | *Neutral* | *Negative* | *Neutral* | *Negative* |
| **Valence** | 5.29 (0.31) | 2.64 (0.70) | 5.11 (0.33) | 2.97 (0.87) |
| **Arousal** | 2.62 (1.51) | 6.13 (1.45) | 2.54 (1.58) | 5.70 (1.38) |
|  | **Words** |  |  |  |
|  | *Attend* |  | *Distancing* |  |
|  | *Neutral* | *Negative* | *Neutral* | *Negative* |
| **Valence** | 5.30 (0.44) | 3.08 (0.55) | 5.11 (0.30) | 3.43 (0.74) |
| **Arousal** | 2.95 (1.56) | 5.06 (1.49) | 2.77 (1.50) | 4.33 (1.64) |

We also computed the pure effect of Regulation (collapsing for other relevant conditions). For both Arousal and Valence, distancing had a significant effect on SAM ratings when compared to attend (respectively, p<0.05; p<0.001). In other words, distancing reduced the arousal and increased the valence of stimuli.

For Words, analyses returned a significant effect of Regulation (F(1,29)=16.738, p <0.001), an interaction between Index and Regulation (F(1,29)=17.606, p <0.001), Index and Emotional content (F(1,29)=262.059, p <0.001), and the triple interaction Index, Regulation and Emotional content (F(1,29)=20.248, p <0.001). To explore the triple interaction, we computed Bonferroni corrected post hoc (threshold p<0.02) on the arousal and the valence of each category. For Arousal, there was a significant difference between distancing and attend for negative words (p<0.001), but not for neutral words (p=0.044). Distancing decreased the arousal of negative words. Neutral and negative words differed from each other in both conditions (p<0.001). See Table 1 and Figure 1B. For Valence, there was a significant difference between distancing and attend for negative words (p<0.005), but not for neutral words (p=0.014). In other words, distancing increased the valence of negative words. Neutral and negative words differed from each other in both conditions (p<0.001). See Table 1 and Figure 1B. We also computed the pure effect of Regulation (collapsing for other relevant conditions). For both Arousal and Valence, distancing had a significant effect on SAM ratings when compared to attend (respectively, p<0.001; p<0.001). In other words, distancing reduced the arousal and increased the valence of stimuli.

### ERPs results

ERPs results were separated into Effects of Strategy (distancing versus attend conditions during the time window of the presentation of the strategy), and Effects of Regulation (distancing vs attend when stimuli are presented).

### Effect of strategy

Visual inspection of the targets grand-averages shows that, compared to the attend condition, the regulate condition elicited a broadly distributed centro-frontal negativity along with a posterior positivity. The difference appears for both picture and word targets, although it seems more pronounced for pictures. For its polarity and distribution, the effect may be classified as a Stimulus Preceding Negativity (SPN), a slow and long-lasting component usually occurring in anticipation of a variety of cognitive and emotional task-relevant stimuli [e.g., 30]. To investigate whether the difference between the attend and distancing condition was significant, analyses were run in four consecutive time windows of 500 ms each, starting 500 ms after strategy presentation and ending and the target presentation. Following the visual inspection of grand-averages, a fronto-central pool of electrodes was created by averaging the channels: Fpz, F1, F2, Fz, FC1, FC2, FCz, Cz, CPz. A 2 x 2 ANOVA with Strategy (attend vs distancing) and Target (pictures vs words) as within-participants factor was run.

In the first three time-windows, no effect reached significance (1000-1500ms time window: all Fs < 2.6, ps >.1; 1500-2000ms time window: all Fs < 3.1, ps >.09; 2000-2500 ms time window: all Fs < 2.9, ps >.1). In the fourth time-window, i.e., between 2500 and 3000 ms after stimulus onset, there was a main effect of Strategy (F (1, 29) = 4.49, p = .04), with a larger negativity for the regulate (MEAN = -.95μV) than the attend condition (MEAN: -.32μV); no further effect reached significance (Target: F <1; Strategy x Target: F <1). See Figure 2A, B, C, D.

**Figure 2.**
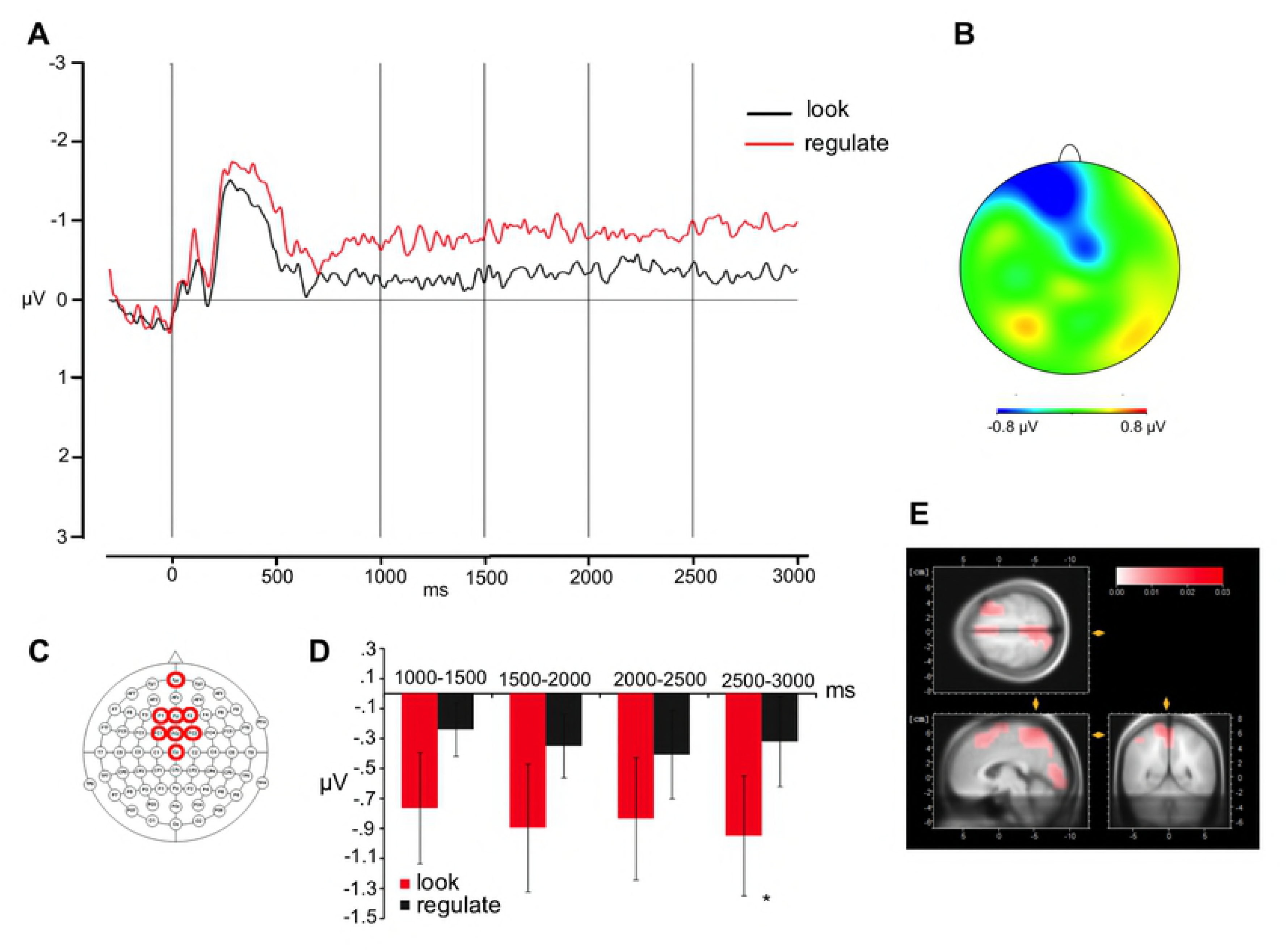
Effect of strategy. Grand average waveforms (A) at the frontal sites (indicated in C) and topographical distribution of the effect distancing minus attend (B); the first vertical line indicates the target onset; the following four vertical lines indicate the boundaries of the time windows for the analyses (1000, 1500, 2000, and 2500 ms, respectively); average activity within each time window is shown in (D). Between 2500 and 3000 ms after stimulus onset, there was a main effect of Strategy with a larger negativity for the distancing than the attend condition. (E) LORETA source reconstruction showed as the maximal source the occipital gyrus, the right middle frontal gyrus, the superior frontal gyrus, the precentral and postcentral gyrus, and the precuneus. Error bars indicate the standard error of the mean, asterisks significant differences.

In order to visualize the neural generators of these scalp effects, and make eventual comparison with previous fMRI experiments, low-resolution brain electromagnetic tomography (LORETA [31]) was performed. This method has been widely described and used in the literature; here we use it in order to obtain likely source estimations for the time window in which significant effects were reported. LORETA source estimates were generated by means of BrainVision Analyzer 2 (for a similar approach, see, e.g., [32], with the current density values provided for 2394 voxels in the gray matter and the hippocampus of a reference brain (MNI 305, Brain Imaging Centre, Montreal Neurologic Institute). As shown in Figure 2C-D-E, the estimation for the comparison between the regulate and the attend condition in the 2500-3000 ms time window showed as the maximal source the occipital gyrus, the right middle frontal gyrus, the superior frontal gyrus, the precentral and postcentral gyrus, and the precuneus. Notably, these areas partially overlap with regions found to be active during regulation tasks in previous fMRI studies [3,15,16].

### Effects of regulation on pictures and words

Pictures and words elicit quite different electrophysiological patterns, both in terms of evoked component and response amplitude. Therefore, the two types of target were analyzed separately. In both cases, two analyses were run: First, we compared the unpleasant and neutral stimuli in the attend-strategy condition in order to identify the electrophysiological response to emotional stimuli. Second, we tested whether the application of a strategy affected the emotional response. To do that, we compared differential waves of targets in the two strategies; specifically, for each strategy the waves of unpleasant targets were subtracted to those of neutral targets and the comparison occurred between subtracted waves. Such analyses were run using the same electrodes groups and the same time windows adopted in the first analysis in which we estimated the emotional effects for the targets in the attend condition only.

## Pictures

### Effect of content for pictures

#### Early Negativity

The inspection of grand-averages shows that, at the occipital sites, unpleasant pictures differ very early from neutral pictures, with the former being more negative than the latter. For its distribution and time dynamic, this negativity resembles a N170. Previous studies reported an emotional modulation of N170 component for emotional facial expressions [33,34] as well as for emotional pictures [35]. Another possible interpretation of this negativity might be in terms of Early Posterior Negativity (EPN). Although its time dynamic (before 200 ms) is a bit faster than that usually associated with such component, a similar time dynamic has been previously reported [36]. The statistical analysis conducted in the 100-170ms time window on a group of occipital channels (O1, Oz, O2) shows a maximal pronounced effect. A one-way ANOVA with the Emotional content (negative vs neutral) of the stimulus as within-participant factor showed the effect to be significant (F (1,29) = 6.40, p =.01), with negative stimuli (1.4μV) being less positive than neutral ones (2.2 μV). See Figure 3A.

**Figure 3.**
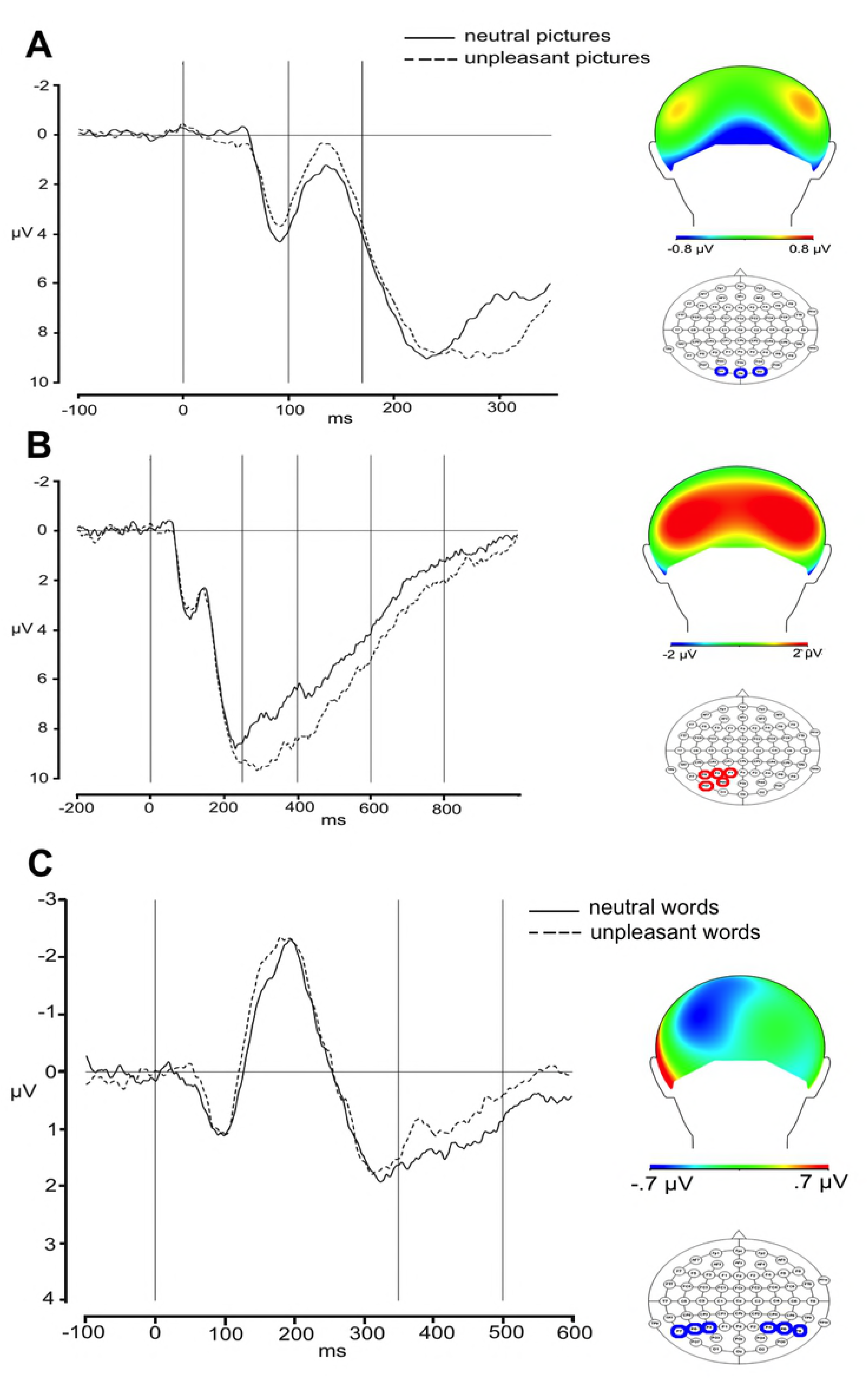
Effect of emotion for pictures. Grand average waveforms for relevant sites and topographical distribution of the emotional effect (negative minus neutral) for pictures (A, B) and words (C). In each figure, the first vertical line in the plot of grand averages indicates the target onset, and the following vertical lines the boundaries of the time windows of analyses.

#### Late Positivity

Negative pictures show a larger positivity on the posterior sites and a corresponding negativity on the centro-frontal area. The two conditions start to differ around 250 ms after target presentation and the positivity appears more accentuated on the left-posterior sites, lasting until approximately 800ms.

To test the statistical significance of the effects, six groups of electrodes were selected (Frontal Left: F5, F3, F1, FC5, FC3, FC1; Frontal Right: F6, F4, F2, FC6, FC4, FC2; Central Left: T7, C3, C1, CP5, CP3, CP1; Central Right: T8, C4, C2, CP6, CP4, CP2; Posterior Left: P5, P3, P1, PO7, PO3, O1; Posterior Right: P6, P4, P2, PO8, PO4, O2). The groups were created on the basis of the grand average waveforms reflecting the topography and temporal dynamics of the brain potential. Statistical analyses were run on 3 consecutive time windows. In all cases, we ran a 2 x 3 x 2 ANOVA with Emotional content (negative vs. neutral), Anterior-Posterior (frontal vs central vs posterior electrodes), and Hemisphere (left vs right) as within-participant factors. Central line was analyzed separately in a 2 (Emotional content: negative vs. neutral) x 3 (Anterior-Posterior: frontal [Fpz, Fz] vs. central [FCz, Cz, CPz] vs. posterior electrodes [Pz, Oz]) ANOVA, with both factors as within-participants. The Geisser and Greenhouse [37] correction was applied to all repeated measures with more than one degree of freedom (only corrected p values are reported).

#### First time window – 250ms-400ms

The main effect of Emotional content was significant (F (1,29) = 47.94, p <.001) as well as the Emotional content by Anterior-Posterior interaction (F(2,58) = 38.78, p <.001). Planned comparison showed that a difference between negative and neutral pictures only emerged on the posterior sites (negative: 9.4 μV vs neutral: 7.6 μV, t = -2.00, p =.04; frontal: negative: -6.4 μV vs. neutral: -5.4 μV, t = 1.64, p >.1; central: negative: 0.1 μV vs. neutral: 0.1μV, t <1; in this and the following planned comparisons p values were adjusted with Bonferroni method). The main effect of Anterior-Posterior was also significant (F(2,58)=99.75, p <.001). No further effect reached significance (Hemisphere: F = 1.77, p >.1; Anterior-Posterior x Hemisphere: F = 1.65, p >.2; all other Fs<1). See Figure 3B.

The analysis of central line showed a significant main effect of Anterior-Posterior (F(2,58)=52.21, p <.001); Anterior-Posterior also interacted with Emotional content (F(2,58)=8.79, p =.001). Planned comparison did not reveal any significant contrast (all ts <1; frontal: negative: -7.2 μV vs neutral: -6.0 μV; central: negative: -3.9 μV vs neutral: -3.4 μV; posterior: negative: 5.2 μV vs neutral: 4.3 μV,).

#### Second time window – 400ms-600ms

Again, the main effect of Emotional content was significant (F (1,29) = 58.11, p <.001) as well as its interaction with Anterior-Posterior (F (2,58) = 6.43, p =.003). Planned comparison, however, did not reveal any significant difference (posterior: negative: 6.8 μV vs neutral: 5.7 μV; t = -1.50, p >.1, central: negative: 1.5 μV vs neutral: 1.1 μV; t = - 1.58, p >.1; frontal: negative: -5.1 μV vs neutral: -4.6 μV; t <1). The main effect of Longitude was significant (F(2,58)=91.32, p <.001). No further effect reached significance (Hemisphere: F = 2.61, p >.1; Emotional content x Anterior-Posterior: F = 1.62, p >.2; Anterior-Posterior x Hemisphere: F = 1.91, p >.1; all other Fs<1).

In the analysis of the central line, the effect of Anterior-Posterior was significant (F(2,58)=35.02, p <.001). No further effect reached significance (Emotional content: F <1; Emotional content x Anterior-Posterior: F = 1.50, p >.2). See Figure 3C

#### Third time window – 600ms-800ms

The main effect of Emotional content was significant (F (1,29) = 33.61, p <.001), as well as its interaction with Hemisphere (F (1,29)=7.95, p =.008). Also the three-way interaction among Emotional content, Anterior-Posterior, and Hemisphere approached significance (F (2,58) = 3.19, p =.07). Planned comparisons showed that negative and neutral pictures differed at left central sites (negative: 1.5 μV vs neutral: 0.5 μV, t = -2.49, p =.01; no further comparison reached significance). The main effect of Anterior-Posterior was also significant (F (2,58)=31.06, p <.001), as well as its interaction with Hemisphere (F(2,58)=4.72, p =.02). No further effect reached significance (Hemisphere: F = 2.8, p >.1, Emotional content x Hemisphere F <1).

In the analysis of the central line, the effect of Anterior-Posterior was significant (F(2,58)=11.10, p <.001). No further effect reached significance (Emotional content: F <1; Emotional content x Anterior-Posterior: F = 2.22, p >.1).

### Effect of regulation on pictures

#### Early Negativity

The one-way ANOVA with Regulation (distancing vs attend) as a within-participant factor showed no significant effect (F <1). To further ascertain that the application of strategy does not modulate the effect we reported (effect of regulation), a 2 x 2 ANOVA on un-subtracted waves was run with Regulation (distancing vs attend) and Emotional content (negative vs. neutral) as within-participants factor. The main effect of Emotional content was significant (F(1,29) = 9.63, p =.004). No further effect reached significance (Regulation: F =3.86, p >.05; Regulation x Emotional content: F <1). The absence of an interaction, together with the absence of any effect in the analysis of the differential waves, clearly indicates that the early responses elicited by negative pictures cannot be regulated through the application of an explicit strategy.

### Late Positivity

For each time window a 2 x 3 x 2 ANOVA was run with Regulation (distancing vs attend), Anterior-Posterior (frontal vs. central vs posterior) and Hemisphere (left vs right) as within-participant factors. The central line was analyzed separately in a 2 (negative vs neutral) x 3 (frontal vs. central vs. posterior) ANOVA, with both factors as within-participants.

#### First time window – 250ms-400ms

The main effect of Regulation approached significance (F(1,29)=3.19, p =.08). Regulation significantly interacted with Anterior-Posterior (F(2,58) = 5.78, p =.001). The three way interaction was also significant (F(2,58) = 5.12, p =.002). Planned comparisons showed differences in the posterior left (distancing: 0.79 μV vs attend: 1.74 μV, t=2.12, p =.04) and frontal right sites (distancing: 0.14 μV vs attend: -1.12 μV, t = -2.52, p =.01). The main effect of Anterior-Posterior was significant (F(2,58) = 18.71, p <.001), whereas Hemisphere approached significance (F(1,29)=4.06, p =.06). No further effect reached significance (Regulation x Hemisphere: F = 1.83, p >.1; Anterior-Posterior x Hemisphere: F = 1.36, p>.2).

The analysis of the central line shown main effects of Regulation (F(1,29) = 6.68, p =.01) and Anterior-Posterior (F(2,58) = 4.07, p =.03). The Regulation by Anterior-Posterior interaction was also significant (F(2,58) = 6.18, p =.008). Planned comparisons showed that the attend condition was more negative that the distancing condition in the frontal site (distancing: -0.06 μV vs attend: -1.18 μV, t = -2.11, p =.04); also central sites approached significance (t = -1.8, p >.07). No difference emerged posteriorly (t =1.26, p >.2).

#### Second time window – 400ms-600ms

The main effect of Regulation approached significance (F(1,29) = 3.32, p =.08), as well as the two-way interaction between Regulation and Hemisphere (F(1,29) = 3.24, p =.08), and the three-way interaction among Regulation, Anterior-Posterior, and Hemisphere (F(2,58) = 3.05, p =.07). The Regulation by Anterior-Posterior was significant (F(2,58) = 6.52, p =.01). Planned comparisons for all the sites shown that the distancing and the attend condition differed at posterior left (distancing: 0.21 μV vs attend: 1.36 μV, t = 2.53, p =.04) and frontal right sites (distancing: 0.56 μV vs attend: -0.73 μV, t = -2.61, p =.01). The other effects were not significant (Anterior-Posterior: F = 2.32, p >.1; Anterior-Posterior x Hemisphere: F = 1.48, p >.2).

The analysis of the central line shown a main effect of Regulation (F(1,29) = 9.87, p =.003) and a Regulation by Anterior-Posterior interaction (F(2,58) = 5.001, p =.01). Planned comparisons showed that the attend condition was more negative than the distancing condition in the central site (distancing: 0.37 μV vs attend: 1.28 μV, t = -2.08, p =.04); also frontal sites approached significance (distancing: -0.78 μV vs attend: 0.40 μV, t = -1.76, p =.08). No difference emerged posteriorly (t =1.02, p >.3).

#### Third time window – 600ms-800ms

The Regulation by Hemisphere interaction approached significance (F (1,29) = 3.56, p =.06). The three-way interaction among Regulation, Anterior-Posterior, and Hemisphere was significant (F (2,58) = 5.72, p =.01). Planned comparisons reveled that, at this point, the distancing and attend condition only differed at the frontal right site (attend 0.38 μV vs attend: -0.91 μV,t = -2.35, p =.02). In the analysis, no further effect reached significance (Regulation x Anterior-Posterior: F = 1.48, p >.2; all other Fs <1).

The analysis of the central line showed a main effect of Anterior-Posterior (F(1,29) = 5.26, p =.02). No further effect reached significance (Regulation: F = 2.51, p >.1, Regulation x Anterior-Posterior: F <1). See Figure 4A, B, C.

**Figure 4.**
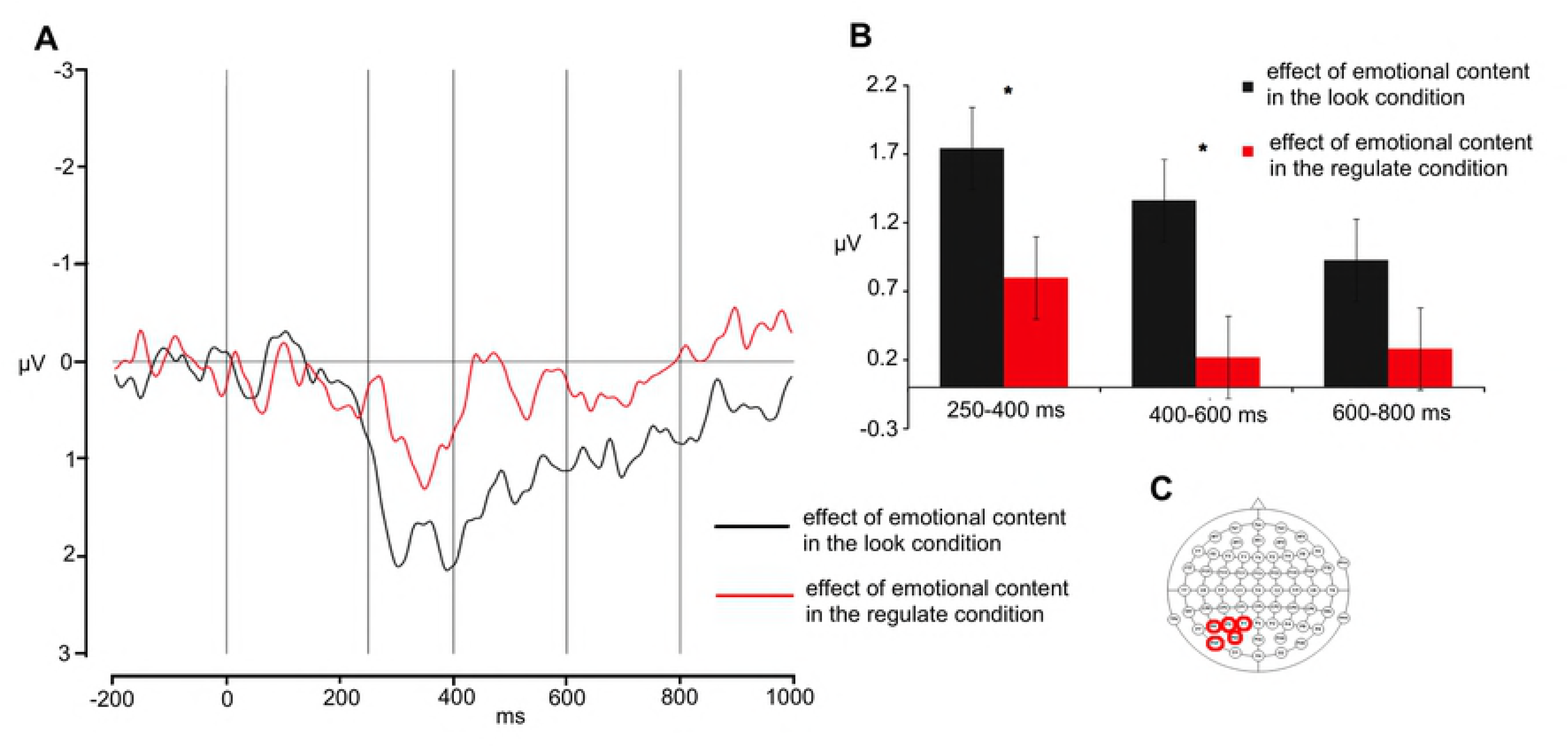
Effect of regulation for pictures. Grand average waveforms (A) at relevant sites (indicated in C) of the effect of regulation for pictures. The first vertical line indicates the target onset; the following four vertical lines indicate the boundaries of the time windows for the analyses (250, 400, 600, and 800 ms, respectively); average activity within each time window is shown in (B). Error bars indicate the standard error of the mean, asterisks significant differences.

## Words

### Effect of content for words

#### Early negativity

The visual inspection of grand-averages seems to show tiny differences between emotional and neutral stimuli. A larger negativity for emotional than neutral stimuli is visible on the parieto-occipital sites starting approximately 350 ms after target onset. For its distribution and polarity, such effect resembles an EPN (see [38] for similar results). To test its statistical significance, we run an analysis on two groups of electrodes (posterior-left: P7, P5, P3, O1; posterior-right: P8, P6, P4, O2), which we selected on the basis of the grand average waveforms reflecting the topography and temporal dynamics of the brain potential (for a similar selection, cfr. [38]). A 2 x 2 ANOVA with Emotional content (negative vs neutral) and Hemisphere (left vs. right) as within-participants factor was run on mean EEG amplitudes in the 350ms-500ms time window. The analysis showed a main effect of emotional content, with negative words producing more negativity than neutral words (1.71 μV vs 2.17 μV, F (1,29) = 4.23, p =.04). No further effect reached significance (both Fs <1).

#### Late Positivity

The electrophysiological data do not show any larger positivity for emotional than neutral stimuli, showing no evidence for the LPP in any area of the scalp. To ascertain the absence of the LPP, statistical analyses were run on three groups of electrodes – i.e., left (CP3, P1, P5, PO3), central (CPz, Pz), and right (CP3, P1, P5, PO3) – on which, according to the literature, the LPP is maximal (e.g., Citron, 2012; Herbert et al., 2008). The 2 (Emotional content: negative vs neutral) x 3 (Topography: left vs. central vs. right) ANOVA on the 500-700 ms time window showed no significant effect (Topography: F = 2.96, p >.06; all other Fs < 1, ps >.5). See Figure 3.

### Effect of regulation on words

The 2 (Regulation: distancing vs attend) x 2 (Hemisphere: left vs. right) ANOVA showed no significant effect (Regulation: F <1; Regulation x Hemisphere: F = 3.2 p>.08; Hemisphere: F = 1.93, p >.1). The result was further confirmed by the 2 (Regulation: distancing vs attend) x 2 (Emotional content: negative vs neutral) x 2 (Hemisphere: left vs. right) ANOVA we ran on un-subtracted waves, that showed only a main effect of Emotional content (F (1,29) = 5.40, p =.02) (Emotional content x Hemisphere: F = 2.16, p >.1, all other Fs <1). The absence of any effect of regulation suggests that the application of a regulation strategy does not have any effect on the first electrophysiological response to emotional written stimuli.

## Discussion

This study provides behavioral and physiological evidence on the possibility to regulate emotional pictures and words stimuli by distancing. At the behavioral level, all participants were able to regulate the negative stimuli presented. Participants were able to regulate both the arousal and the valence of negative pictures in such a way as to reduce their emotional impact (decreased arousal, effect magnitude of [Distancing_(negative pictures)_ - Attend_(negative pictures)_] = -0.42 points), and to decrease their negativity (increased valence, +0.32)). While the regulation of pictures is in line with previous findings and confirms them [17,39-41], the novelty of the study is the demonstration that negative words can be regulated in both the arousal and the valence dimensions by using the strategy of distancing. Arousal decreased (effect magnitude of [Distancing_(negative words)_ - Attend_(negative words)_] = -0.73 points) and valence increased (+0.35) as an effect of strategy. To our knowledge this is the first time that emotional words are shown to be regulated in both arousal and valence by the use of distancing. This finding has some relevant implications. First, we constantly use words to regulate our emotions that are elicited when talking with others; second, we may want to regulate others emotions by using words. Consider the example of what happens during social interactions. Understanding how we regulate linguistic stimuli is thus of fundamental importance in order to understand and possibly maximize how to regulate emotions. Moreover, previous studies rarely took into account both arousal and valence (see for an exception [15], and [22]). This is especially true for the case of words. We showed separate behavioral effects of regulation on both arousal and valence.

A second finding is that distancing can be indeed an effective strategy for regulating both words and pictures and both emotional dimensions comparable to other well studied strategies such as reappraisal or distraction. This result is important for two reasons. First, distancing is a commonly used strategy that does not require any particular cognitive effort (reinterpreting the meaning of the stimulus, or engaging in a distracting task). Clinicians have long reported the use of distancing-like strategies (e.g. isolation of affect, experiential avoidance, behavioral avoidance, dissociation, and others) to protect ourselves from unbearable life events we cannot avoid or escape from. Second, from a methodological point of view, having shown that distancing is effective in regulating emotions, opens the possibility of using such strategy in studies in which more complex strategies may be difficult to use. Also, the fact that distancing is effective for both pictures and words implicates that can be successfully used in different situations independently of the nature of the emotional stimuli (linguistic, pictorial, etc). This may also indicate that this strategy is modality independent and that it may act by blocking the emotional impact of stimuli without affecting the meaning of the stimuli themselves. This is very different from other strategies (e.g. reappraisal) that change the meaning of the stimuli in order to affect their emotional impact. Future studies may want to compare these strategies to understand more exactly what aspect they alter in the sequence of elaboration of the stimuli.

In this study, ERPs were recorded to identify neural markers along the regulation process. Before analyzing the signal associated with the regulation of pictures and words, we analyzed the signal relative to the implementation of the strategy (in the time window when strategies are probed) as the first temporal event occurring during each trial. Notably, only few experiments explored the effect of strategy [23-25], as they commonly focused on the effect of regulation (during stimulus presentation). We found that when participants applied the distancing strategy there was a significant long-lasting negativity over frontocentral electrodes starting approximately 2.5 s after the strategy presentation and lasting until the target appeared: the negativity was larger for the regulate than for the attend condition and was not modulated by the to-be-regulated stimulus, i.e. it was equal for both pictures and words. We interpreted the effect as belonging to the class of components known as Stimulus-Preceding Negativity (SPN), usually occurring during the warned fore period preceding a task (regulate the response of the following emotional stimuli in our case). SPN has been reported by previous experiments investigating the electrophysiological response to strategy implementation [23,25], and has been interpreted as reflecting enhanced attentional orienting to and anticipation of impeding stimuli This negativity may be also related to what a previous fMRI experiment found when analyzing the effect of strategy (more activation compared to the no strategy condition) (see [15,16]). Previous studies showed that the regulation of unpleasant stimuli, may be related to the activity of frontal regions (e.g. lateral and medial frontal gyri, pre-supplementary motor area, and cingulate cortex) (see [15,16]). Dipole reconstruction confirmed the source of this effect may involve large portion of the lateral and medial frontal cortex. This is in line with previous neuroimaging findings suggesting that down-regulation of unpleasant emotions activate prefrontal and cingulate regions (for a review, see [3]). We also found portions of the precuneus and of the occipital lobe not reported in previous fMRI studies. Future studies may want to better separate the role of these regions. Building on our and previous findings, we suggest that beside the type of cognitive strategy to use, a circuit involving the lateral and medial portions of the frontal cortex (but also extending to medial parietal and occipital regions), are responsible for the selection and implementation of the strategy (effect of strategy).

In order to assess possible differences between types of stimuli, the effect of regulation was also investigated through separate analyses for pictures and words. As expected, in the attend condition, pictures showed early and late modulations previously observed in study with emotional stimuli. Unpleasant pictures elicited an early negativity within 200 ms and maximal at occipital sites, with a larger negativity for unpleasant than pleasant stimuli; this component was followed by a larger positivity (i.e., LPP) for negative than neutral stimuli starting ∼250 ms after target onset and lasting until ∼800 ms after target onset; the positivity was clearly visible on the posterior sites and had a corresponding negativity on the centro-frontal area. The pattern of results is coherent with previous findings (e.g., 35,42).

A potential interpretation of our early negative component is in terms of N170, which has been reported to be sensitive to the pictures emotional content [35,43]. In line with a more general interpretation of early negativities in the emotional literature, a modulation of N170 and has been interpreted in relation to a selective attention mechanism that is involved in the detection of salient emotional stimuli to pay attention to for further processing [e.g., 35,43-45]. The early negativity, however, might be also interpreted in terms of EPN. Usually, this component has a slightly slower time dynamic, being visible ∼200-300 ms after target onset, but an early EPN starting 150 ms after target onset has been also reported by Junghofer et al. [36]; note that these authors used a shorter presentation of pictures that might have anticipated the EPN). The EPN has been interpreted as indexing the selective attention, with attention orientation, which is highly sensitive to the stimulus emotionality [35-45]. This early activity might reflect rapid amygdala processing of unpleasant information and increased engagement of autonomic arousal towards salient stimuli [46-48]

Our study showed that negative pictures modulated the early negative components, and that these component is not modulated by the regulation strategy. This may be due to the attentive nature of this component that is necessarily elicited by emotional stimuli, nor subject to cognitive control neither to eventual modulation by the strategy. Previous experiments clarified that early components may be task-independent, as their effect is not modulated by the depth of processing [49], the emotional nature of the task [50], or the self-referentiality of the emotional stimulus [51]. This suggests that these components index automatic, implicit processing of emotion [52]. Scholars agree on the fact that emotion regulation involves a process of early emotional reactivity (mainly automatic), as well as later voluntary inhibitory control [53].

The LPP is typically increased by emotional compared to neutral visual stimuli [19,44,54,55] and reflects enhanced processing and attention to emotional salient stimuli [54]. Larger LPP amplitudes are also correlated with increased arousal [54]. Several studies showed that the LPP can be modulated by reappraisal, showing larger deflections when up-regulating [23] and reduced deflections when down-regulating the emotional response [12,19,22,23,55-57]. The neural generators of the LPP are thought to be the extrastriate visual system and emotion-related structures such as amygdala [58], and may reflect stronger functional connectivity between occipital cortex and frontal areas [59]. Our study confirms this modulation of LPP as an effect of down-regulating the content of emotional pictures (significant difference in the central line electrodes), especially in the first stages of the component (i.e., between 250 ms and 600 ms after target presentation), that showed the maximal LPP reduction at multiple sites. Thus, when comparing the impact of the emotional content on the stimuli in the distancing vs. attend condition, one may immediately note the drastic LPP reduction (left-posterior region, first time window [250 400 ms]: effect of emotional content (negative - neutral) in the REGULATE condition: 0.79 μV vs. effect of emotional content (negative - neutral) in the Attend condition: 1.74 μV; second time window [400 600 ms]: effect of emotional content (negative - neutral) in the REGULATE condition: 0.21 μV vs. effect of emotional content (negative - neutral) in the Attend condition: 1.36 μV). Previous studies using ERPs reported that the LPP amplitude is affected by instructions to down-regulate the emotional responses due to negative stimuli [12,20,21,55,56,60]. These studies involved participants using cognitive strategies such as reappraisal, suppression and distraction, and showed regulation of both subjective ratings and LPP. Our study is the first to show an effect of modulation over LPP due to a not commonly studied strategy defined as distancing. Although distancing was originally proposed by Gross [53] to be a cognitive strategy, we believe that it should be considered not a pure cognitive strategy (no explicit thought generation, nor rethinking effort), but rather an experiential strategy as it involves a different way of relating to the stimuli (see also [1,2,61]). Research has shown that adopting a mindful, or unemotional self-observation attitude, can be a useful mean to reduce emotional reactivity to negative information comparable to, or even more efficient than, cognitive strategies [61-63].

Unpleasant linguistic stimuli elicited a larger negativity for negative than for neutral stimuli; for its timing and occipital distribution, we interpreted such negativity as an EPN. The EPN for emotional words has been repeatedly reported across different tasks: silent reading [38,64,65], lexical decision [49,52,66], written word identification [67] and Stroop [68]. Our study confirms previous findings and is in line with the view that the EPN is task-independent and may be elicited by the simple passive exposure to written material. Moreover, the failure to find any modulation of such component parallels our results with pictures and suggests that the EPN indexes early automatic processing of emotion (e.g., 52, 42). A second possible explanation for the absence of a modulation of the EPN may be that the EPN is a smaller component (circa 0.80 mV) as compared to LPP (1.80mV). This reduced size may have clouded the effect of regulation.

Word stimuli, however, did not show any LPP. The absence of such component may be due to our experimental design. The literature on emotional words shows mixed and contrasting finding on the LPP: while studies usually reported larger LPP for emotional than neutral words [e.g., 67,69,70,49], other studies reported the opposite pattern [e.g., 71], and others no difference at all between emotional and neutral words [67]. Indeed, LPP amplitude is modulated by different dimensions as type of task – with larger amplitude when a more in-depth processing of the stimulus is required [49] – and the emotional self-relevance of the content – with larger amplitude when stimuli are referred to the self rather than to others [e.g., 51]. A further reason for the absence of the LPP with word stimuli may be due to the lesser power words have to induce emotional responses. In a previous experiment, Kensinger and Schacter [72] using fMRI, reported lower statistical power and therefore more circumscribed activations for words than pictures. No effect of modulation for LPP was visible. The modulation of LPP when regulating linguistic stimuli was observed only by Deveney and Pizzagalli [6] probably due to the nature of the task used that combined visual and linguistic stimuli in sequence.

Some limitations of the present study should be acknowledged. First, when words are considered, ERPs data failed to show any regulation effect, representing a dissociation between behavioral and ERPs measures for negative linguistic stimuli. It is not clear whether the lack of ERPs modulation was a limitation of the current task design (selection of stimuli, valence and arousal power, etc), or whether these findings indicate that ERPs might be less sensitive for detecting the effects of regulation over such class of stimuli. We believe the emotional effects of linguistic stimuli are harder to detect. Future studies may focus on linguistic stimuli only by using more arousing and negatively valenced stimuli to increase the possibility of detecting ERPs effects. Although we used the good spatial resolution of 64 electrodes montage, an higher spatial resolution may have help finding finer grained differences.

These limitations notwithstanding, the present study provides strong behavioral evidence that linguistic stimuli can be regulated and this may pave the way to better understand of how our language can be used not only to catch emotional meaning, but also to understand how we regulate emotions conveyed by words. In everyday life we are constantly engaged in social interactions and words are the primary mean we use to express emotional meaning. To prevent excessive interference with ongoing cognitive and social abilities necessary to interact with others, and to avoid being overwhelmed by emotions, incoming linguistic stimuli sometimes must be regulated. This study further supports the possibility that humans may use distancing to regulate emotional pictures and words.

